# TopoFuseNet: Hierarchical Graph Representation Learning with Multi-Scale Topological Features for Accurate Drug Synergy Prediction

**DOI:** 10.64898/2026.05.05.722940

**Authors:** Xiaoyang Shi, Qi Wang

**Affiliations:** College of Science, China Agricultural University, Beijing 100083, China

**Author notes:** Corresponding author. E-mail address (Q. Wang).

**Keywords:** Drug Combination Prediction, Graph Neural Network, Hierarchical Representation, Group Centrality, Multi-modal Fusion

## Abstract

Accurate prediction of drug synergy is paramount for developing effective combination therapies and advancing personalized medicine. Although methods based on graph neural networks (GNNs) have become a prevalent approach, they often treat molecules as flat graphs of connected atoms, thus overlooking their inherent hierarchical structure (i.e., atoms forming functional groups) and the critical topological information that governs molecular interactions. To address this limitation, we introduce TopoFuseNet, a novel hierarchical graph representation learning framework that integrates multi-scale topological features. The core innovations of TopoFuseNet include: 1) The first-ever application of “Group Centrality” from network science to cheminformatics, enabling the identification and quantification of functional groups crucial to drug activity; 2) A systematic, multi-path strategy to seamlessly integrate node-level (atom) and group-level (functional group) topological features into a Graph Attention Network (GAT) via feature augmentation, attention biasing, and hierarchical pooling; 3) A Differential Transformer module to deeply fuse multi-modal features learned from sequences, fingerprints, and our proposed hierarchical graph representations.

Extensive experiments on two large-scale benchmark datasets, DrugComb and DrugCombDB, demonstrate that TopoFuseNet significantly outperforms state-of-the-art methods across multiple key metrics, including AUC, AUPRC, and F1-score, while exhibiting exceptional generalization robustness under various stringent cold-start scenarios. In-depth ablation studies further confirm the effectiveness and necessity of each proposed innovative module. Furthermore, multi-scale interpretability analysis and zero-shot cross-domain transfer experiments reveal that the model successfully captures molecular interaction rules with clear pharmacological significance, demonstrating immense practical potential for discovering novel combination therapies through large-scale virtual screening. Our work not only delivers a superior model for drug synergy prediction, but more importantly, it establishes a novel and scalable paradigm for effectively integrating hierarchical molecular structures and topological information into GNNs.

## 1. Introduction

Combination therapy, which involves the concurrent use of multiple drugs, has become a core strategy for treating complex diseases such as cancer. Successful drug combinations can produce synergistic effects, where the combined outcome is greater than the sum of the individual drug effects. This leads to enhanced therapeutic efficacy, reduced dosages, and a lower likelihood of developing drug resistance, which are the primary goals of combination therapy research [1-3]. However, the number of potential drug combinations grows combinatorially, making it impractical to screen all possibilities through traditional in vitro or in vivo experiments. This reality makes the development of efficient and accurate computational models for predicting drug synergy particularly urgent. Accurate synergy prediction can not only significantly accelerate the drug discovery and development pipeline and reduce R&D costs but also provide critical decision support for clinicians, help avoid potentially harmful drug combinations, and drive the advancement of personalized medicine [4].

Early computational methods relied primarily on knowledge bases or similarity-based assumptions. Knowledge-based methods employed logical reasoning based on expert-defined pharmacological rules [5], but they suffered from low coverage and were unable to predict novel interactions. Similarity-based methods, on the other hand, assumed that drugs with similar chemical structures or targets would exhibit similar interaction patterns [6,7,47]. These approaches struggle to capture complex non-linear relationships and perform poorly in predicting the effects of new drugs.

In recent years, deep learning, particularly Graph Neural Networks (GNNs), has achieved tremendous success in the field of drug synergy prediction. Pioneering work such as DeepSynergy first introduced deep feed-forward networks, demonstrating the potential of deep learning [8]. Subsequently, models like DeepDDS [9] marked a breakthrough by converting drugs’ SMILES sequences into molecular graphs and using GNNs to learn their structural representations, achieving a significant leap in performance. More recent methods, such as DFFNDDS [10], have enhanced sequence features by fine-tuning language models, while others like HDN-DDI [11] have explored hierarchical molecular graphs to further improve prediction accuracy.

However, despite these significant advancements, a fundamental limitation persists in existing GNN-based models: they tend to treat the molecular graph as a flat, homogeneous structure, failing to fully leverage its intrinsic hierarchical organization and complex topological properties, which are crucial for biological activity. In chemical reality, atoms do not exist in isolation; they form functional groups with specific chemical meanings, and these groups are the key units that determine a molecule’s properties and interaction patterns. Current methods inadequately model this hierarchical structure from atoms to functional groups and overlook the rich topological features that describe node importance and network morphology. This has created an information bottleneck in molecular representation learning, thereby limiting the accuracy and interpretability of predictive models.

To bridge the aforementioned research gap, this paper proposes a novel deep learning framework named TopoFuseNet, whose core objective is to learn a more informative and chemically intuitive hierarchical drug representation by systematically modeling and integrating multi-scale molecular topological information. Unlike previous works, TopoFuseNet conceptualizes the molecular structure as a multi-level system comprising both atomic and functional-group levels, and deeply mines its topological properties. The main contributions of this paper are summarized as follows:

1. Proposing a novel hierarchical graph representation learning paradigm. This paradigm introduces the concept of Group Centrality [12] from network science into drug molecule analysis for the first time. It is utilized to quantify the importance of key functional groups within the molecular structure, thereby enabling the model to focus on the chemical groups that are decisive for drug interactions.
2. Designing a modular, multi-pathway topological feature fusion mechanism. We systematically explore three distinct pathways for integrating atomic-level topological features (e.g., node centrality) and functional-group-level topological features (e.g., group centrality) into the GNN: as a supplement to initial node features, as a dynamic bias for graph attention computation, and as a guiding weight for hierarchical graph pooling. This design offers a high degree of flexibility to maximize the utility of topological information.
3. Constructing a robust multimodal feature fusion and prediction module. TopoFuseNet employs an advanced Differential Transformer architecture capable of capturing and integrating the complex non-linear dependencies among various information sources (BERT-based sequence embeddings, chemical fingerprints, and our proposed hierarchical graph representation), thereby making more accurate synergy predictions.
4. Conducting comprehensive performance evaluations and architectural validations. In addition to significantly outperforming existing SOTA models on two large-scale benchmark datasets, TopoFuseNet consistently demonstrates exceptional robustness across four stringent cold-start scenarios. Furthermore, in-depth ablation studies and parameter sensitivity analyses fully corroborate the necessity of each proposed innovative component and the stability of the model architecture.
5. Comprehensively revealing the model’s representational discriminative power, multi-scale interpretability, and clinical translational value. t-SNE feature clustering confirms that the model has learned highly discriminative latent manifolds; multi-scale attention profiling indicates that the model can not only precisely pinpoint key pharmacophores but also possesses the reasoning capability to dynamically adjust cross-modal attention. Furthermore, based on the outstanding virtual screening potential demonstrated in zero-shot cross-domain scenarios, we successfully discovered novel synergistic combinations supported by literature through an *in silico* screening of nearly 580,000 candidate combinations.

## 2. Methodology

### 2.1. Problem Formulation

In this study, the drug-drug interaction (DDI) prediction task is formulated as a binary classification problem. Given a drug pair(*D*_*i*_, *D*_*j*_)and their corresponding cell line context feature *C*_*k*_, the objective of the model is to learn a mapping function *f*. This function takes the multi-modal feature representations *x*_*i*_ and *x*_*j*_of the two drugs and the cell line feature *C*_*k*_ as inputs, and outputs a probability value *p* ∈ [0,1]. This value represents the likelihood that drugs *D*_*i*_ and *D*_*j*_ will exhibit a synergistic interaction in cell line *C*_*k*_.

This can be expressed as:

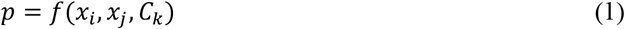

### 2.2. Datasets and Feature Engineering

We utilized two large-scale, widely-used public datasets: DrugComb [13] and DrugCombDB [14], filtering for experiments with corresponding gene expression profiles from the Cancer Cell Line Encyclopedia (CCLE) database. This resulted in a final DrugComb dataset of 292,005 experiments (3038 drugs, 98 cell lines) and a DrugCombDB dataset of 106,709 experiments (1084 drugs, 72 cell lines). For our binary classification task, synergistic interactions were labeled as positive samples and all others as negative.

For each drug pair-cell line triplet, we constructed a multi-modal feature set, as detailed below:

1. Drug Features: Each drug is characterized from three key perspectives:
  a. Sequential Representation: A fine-tuned SentenceTransformer model is used to encode the SMILES string into a 384-dimensional semantic vector.
  b. Substructural Representation: A 1024-dimensional Morgan fingerprint (ECFP) is generated using RDKit to represent the molecule’s chemical substructures.
  c. Graphical Representation: The molecule is converted into a graph, into which three customized features are injected for use by the Graph Neural Network (GNN): 1. Node Feature Augmentation: The 8- dimensional basic chemical attributes of an atom are concatenated with its topological features. 2. Attention Guidance: Independent atomic topological metrics are used as a topological bias in the Graph Attention (GAT) layers. 3. Hierarchical Pooling: The group centrality of functional groups and the atomic membership to these groups are calculated to guide the subsequent hierarchical graph pooling.
2. Biological Context: The features of each cell line are represented by its 18,046-dimensional gene expression profile vector.

Ultimately, the model takes this multi-view feature set, composed of features from Drug A, Drug B, and the specific cell line, as input to predict synergy.

To comprehensively and multi-dimensionally evaluate the generalization capability and robustness of TopoFuseNet, we designed a comprehensive experimental framework comprising a traditional Random Split and four progressive Cold Start splitting scenarios.

First, under the Standard Random Split setting, we partitioned the data into an 80% training set and a 20% test set; the training set was further subdivided into a 75% training subset and a 25% validation subset.

Furthermore, to authentically simulate the stringent challenges encountered in clinical drug discovery, we designed the following four cold-start settings from the perspective of data distribution:

1. Generalized Combination Cold Start: Serving as a universal baseline, this setting solely requires that the drug combinations in the test set have never appeared in the training set. By imposing no additional restrictions on whether the individual drugs are known, it represents a fundamental evaluation state mixing known and unknown drug components.
2. Global Warm Start: Aimed at simulating “drug repurposing” or “new uses for old drugs” scenarios, this setting mandatorily requires that all individual drugs have appeared in the training set, meaning the test set contains only novel combinations of known drugs.
3. Strict Combination Cold Start: This is a strictly controlled transductive setting. While ensuring that the test combinations are entirely novel, this setting imposes a rigorous constraint: every individual drug within the test combinations must have been paired with other drugs in the training set.
4. Drug-Level Cold Start: Serving as the most challenging inductive setting, this scenario requires that the test set contains at least one entirely novel drug that was completely unseen during the training phase.

### 2.3. TopoFuseNet Model Architecture

Our TopoFuseNet framework (Fig. 1) is composed of a multi-modal feature encoding layer and a Differential Transformer fusion and prediction layer, designed to deeply fuse the sequence, fingerprint, and our proposed hierarchical topological graph representations of drugs.

**Figure 1.**
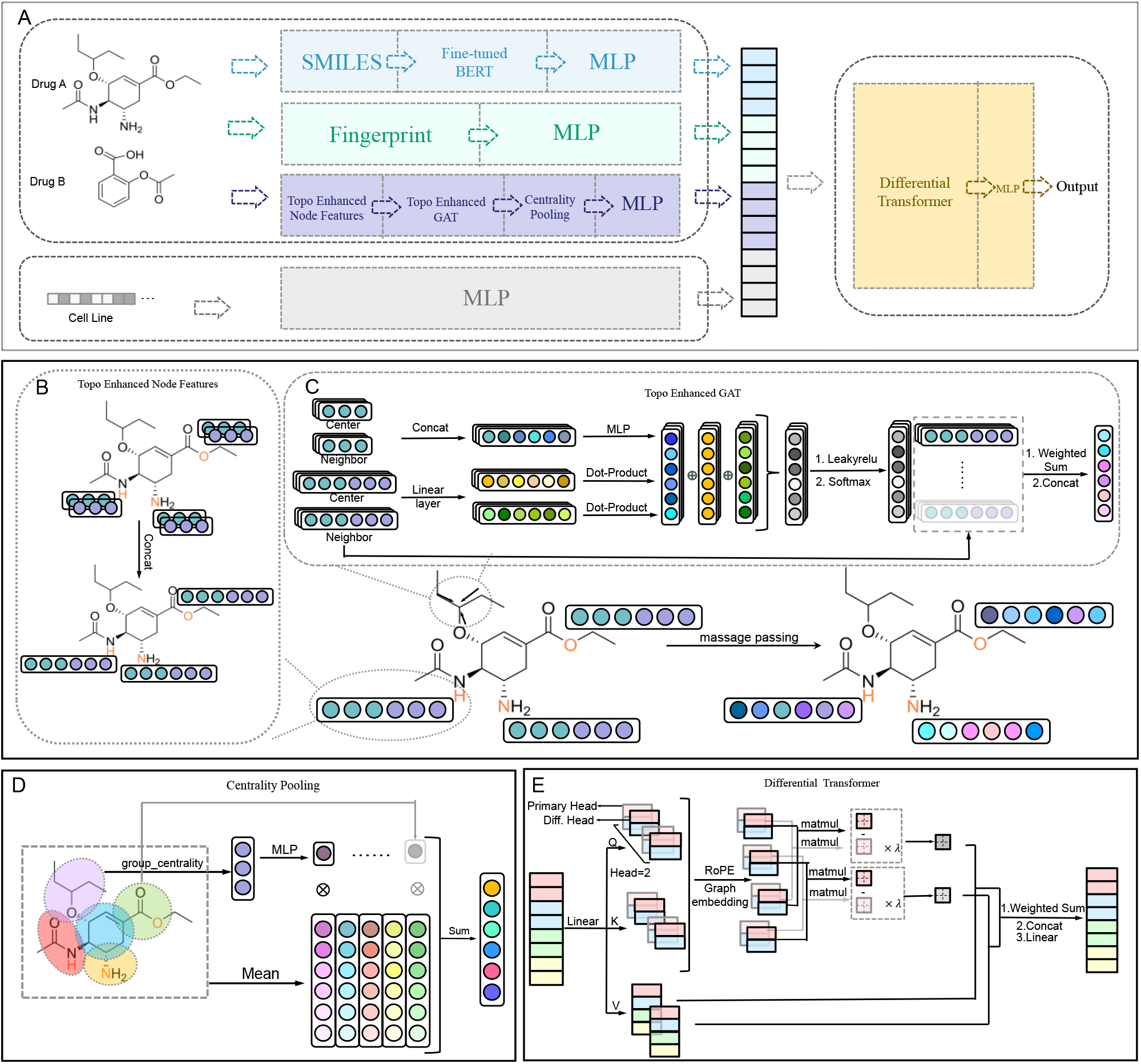
Schematic diagram of the overall architecture and core modules of TopoFuseNet. (A) The multimodal input and feature fusion backbone of the model. Drug molecules extract multi-view features via SMILES, molecular fingerprints, and the topology-augmented graph neural network. After dimensional alignment with cell line features via Multilayer Perceptrons (MLPs), they are jointly fed into the Differential Transformer for cross-modal interaction and final synergy prediction. (B) Topology-Augmented Initial Node Features. The basic chemical attributes of atoms are concatenated with topological attributes to enrich the local structural representations of nodes. (C) Topology-Aware Attention Mechanism (TopoEnhancedGAT). Building upon standard node feature computation, a topological MLP module is introduced to calculate the dynamic topological bias between source and target nodes, guiding the message-passing process with graph-theoretic structural priors. (D) Group Centrality-Guided Hierarchical Pooling. A physically-inspired hierarchical graph pooling method. By extracting specific functional groups within the molecule and calculating their Group Centrality, it generates weights based on group importance, thereby accurately aggregating node-level features into a molecule-level global embedding. (E) Differential Transformer fusion module. Utilizing a Primary Head with Rotary Position Embedding (RoPE) and a Differential Head (Diff. Head), it cancels out common-mode noise across modalities by computing the difference in attention scores, achieving the efficient and denoised fusion of drug and cell line features.

#### 2.3.1. Multi-modal Feature Encoding Layer

This layer consists of four parallel branches that process input information from different sources.

##### 2.3.1.1. Sequential Representation Module

SMILES, as a linearized molecular representation, contains rich structural and syntactic information. To fully exploit this, we selected DeepChem/ChemBERTa-77M-MLM [15, 16] as the base model, which is a BERT model pre-trained on a large database of chemical molecules. We fine-tuned this model using contrastive learning on GuacaMol, a dataset with a vast number of SMILES strings, to enhance the semantic consistency of the generated embeddings. Specifically, we adopted a SimCSE [17] -like strategy: the same SMILES string is passed through the model twice with different dropout masks. The two resulting embeddings are treated as a positive pair, while all other SMILES embeddings within the batch are considered negative samples. We used MultipleNegativesRankingLoss as the loss function for training.

After fine-tuning, this SentenceTransformer model serves as a powerful feature extractor, encoding an input drug SMILES string into a dense, information-rich 384-dimensional embedding vector.

##### 2.3.1.2. Hierarchical Graph Encoder (MolecularGNN)

This is the core innovation of our model. It is a highly modular Graph Attention Network (GAT) engine that systematically integrates multi-scale topological information through the following three pathways:

###### Pathway 1: Topology-Augmented Initial Node Features

The pre-computed feature vector 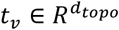, describing the topological attributes of an atom, is concatenated with the basic atomic feature vector 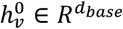 obtained via RDKit to form a more informative enhanced feature vector 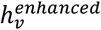. In our model, we specifically adopt Harmonic Centrality as the initial node-level topological feature metric.

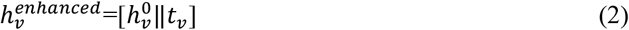

where ‖ denotes the vector concatenation operation. This enhanced vector serves as the input to the first layer of the graph attention network.

###### Pathway 2: Topology-Aware Attention Mechanism

We propose a TopoEnhancedGATConv layer to replace the standard GAT layer, directly injecting topological information into the computation of attention weights. For the edge connecting nodes *i* and *j*, the computation of its attention coefficient *α*_*ij*_ depends not only on the node features but is also modulated by a dynamic topological bias term *b*_*ij*_. This bias term is mapped from the topological features *t*_*i*_ and *t*_*j*_ of the two nodes via a Multi-Layer Perceptron (MLP).

In constructing the topological bias, we primarily introduce Second Order Centrality as the core metric for the node topological feature *t*_*i*_. For an atomic node *i* in a molecular graph *G* = (*V, E*), Second Order Centrality measures the standard deviation of the return times to that node during a Perpetual Random Walk on the graph. From a chemical intuition perspective, introducing Second Order Centrality aims to capture the structural stability and topological correlation of atoms within the molecular skeleton. This mathematical measure of volatility effectively reflects the chemical microenvironment of the atom, enabling the model to a priori perceive and distinguish between the core stable skeleton and active reactive side chains during attention computation. Mathematically, letting *T*_*i*_ be the return time variable for a random walk to node *i*, the Second Order Centrality is defined as:

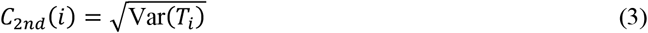

The complete computational workflow is defined as follows:

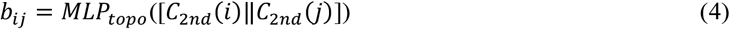

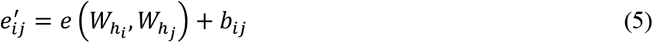

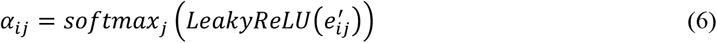

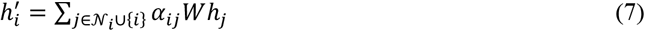

where *e*(·,·) is the standard attention function, *W* is a learnable weight matrix, and 𝒩_*i*_ is the neighborhood of node *i*.

###### Pathway 3: Group Centrality-Guided Hierarchical Pooling

Before graph pooling, the model must first accurately locate key local structures with pharmacological significance within the molecular graph. To this end, relying on classic chemoinformatics rules, we employ a Substructure Matching algorithm based on SMARTS (Smiles Arbitrary Target Specification) patterns to automatically scan and extract the atom sets of various standard functional groups and pharmacophores (such as hydrogen bond donors/acceptors, aromatic ring systems, acidic/basic groups, etc.) contained in the molecule. Through this deterministic rule-based retrieval, the model can reliably map underlying atoms to local groups with explicit chemical characteristics.

Upon completing the group partition, to generate a graph-level representation that captures the core of chemical functions, we designed a GroupAttentionPooling layer. This layer first aggregates node embeddings into a functional group-level representation 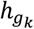. Subsequently, a vector 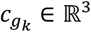 composed of three group centrality metrics for each functional group is utilized to calculate its importance weight *β*_*k*_. The final graph embedding *h*_*G*_ is the weighted sum of all functional group representations.

Specifically, the vector 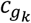 encompasses Group Degree Centrality, Group Closeness Centrality, and Group Betweenness Centrality. For a specific functional group (i.e., a node subset *S ⊂ V*) in the molecular graph *G* = (*V, E*), the rigorous mathematical definitions and chemical intuitions of these three centralities are as follows:

###### Group Degree Centrality

Measures the external connection ratio of a group, reflecting the reaction accessibility and spatial exposure of a pharmacophore within the molecule.

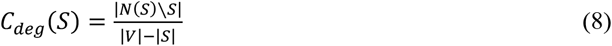

where *N*(*S*) is the set of all nodes directly connected by chemical bonds to the nodes in *S*.

Group Closeness Centrality: Measures the average shortest graph distance from the group to the rest of the molecule, used to evaluate the group’s central hub capacity in transmitting electronic effects such as conjugation and induction.

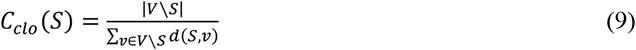

where 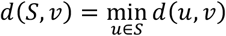 is the shortest distance from node *v* to the atoms within group *S*.

Group Betweenness Centrality: Measures the bridging role played by the group in the shortest paths between other non-group nodes, useful for identifying skeletal structures that support the molecular spatial conformation.

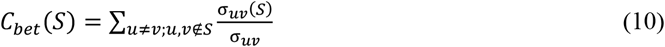

where σ_u*v*_ is the total number of shortest paths between nodes u and *v*, and σ_u*v*_(*S*) is the number of shortest paths passing through group *S*.

For the *k*-th functional group *g*_*k*_ in the molecule, we calculate the three aforementioned group centrality metrics and concatenate them into a three-dimensional group centrality feature vector 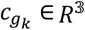:

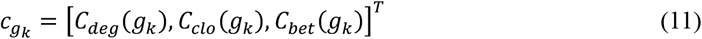

Subsequently, the model utilizes this feature vector 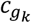to evaluate the global importance of different functional groups, thereby guiding the physically-inspired aggregation of node-level features into a molecule-level global feature. This hierarchical pooling mechanism can be formulated as:

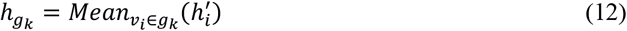

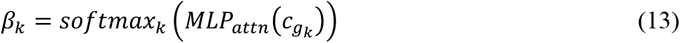

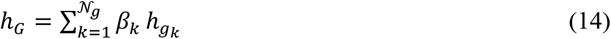

where 𝒩_*g*_ is the total number of functional groups in the molecule, and *MLP*_*attn*_ is a small attention network.

##### 2.3.1.3. Molecular Fingerprint Representation and Cell Line Representation

A 1024-dimensional Morgan fingerprint (ECFP), which serves as a classic and efficient descriptor of chemical structures. An 18,046-dimensional feature vector (gene expression profile) for each cell line, downloaded from the Cancer Cell Line Encyclopedia (CCLE) database[18].

#### 2.3.2. Differential Transformer Fusion and Prediction Layer

Feature vectors from diverse information sources (SMILES embeddings, molecular fingerprints, and graph embeddings output by the GNN), along with cell line features, are first projected into a unified 256-dimensional latent space through their respective independent fully connected layers.

Subsequently, these aligned feature vectors are stacked into a sequence and input into the Differential Transformer [19, 46] fusion module. During multimodal fusion, homogeneous feature redundancy inevitably exists among heterogeneous data. Unlike standard Transformers, which are prone to attention dispersion amidst common-mode noise, the Differential Attention mechanism partitions the attention into paired primary attention heads (subscript 1) and auxiliary attention heads (subscript 2), and introduces Rotary Position Embedding (RoPE) to preserve sequence order. Its core computational formulas are as follows:

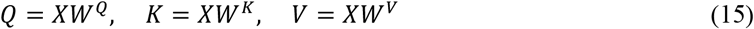

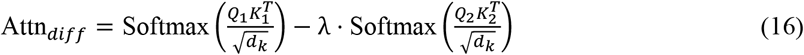

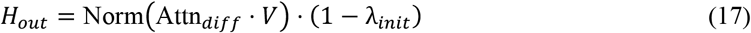

Here, *Q*_1_, *K*_1_, and *Q*_2_, *K*_2_, denote the query and key matrices for the primary and auxiliary heads, respectively, and *d*_*k*_ is the scaling dimension. The scalar λ = exp(λ_*q*1_ · λ_*k*1_ − λ_*q*2_ · λ_*k*2_ + λ_*init*_) is an adaptively learned noise suppression coefficient utilized to dynamically adjust the intensity with which the auxiliary attention head cancels out background noise. Through this subtraction mechanism, the model is capable of acquiring a more sparse, complementary, and discriminative cross-modal interactive representation. Finally, the feature sequence, following deep fusion by the Transformer, is flattened and passed through an MLP classifier to output the probability of the drug combination exhibiting a synergistic effect.

### 2.4. Model Training and Optimization

To effectively train our proposed model and realize its full potential, we designed a systematic training and optimization strategy, covering the loss function, optimizer, regularization, and hyperparameter tuning.

1. Loss Function: For the binary classification task of drug synergy prediction, we employ the Binary Cross-Entropy (BCE) loss as the fundamental training objective. To enhance the model’s robustness and generalization capability, we incorporated the R-Drop regularization strategy. This strategy involves performing two forward passes for the same input batch, each with a different dropout mask. In addition to the BCE loss, a symmetric Kullback-Leibler (KL) divergence loss term is added to penalize the inconsistency between the two output distributions. Specifically, for a given single input sample, we perform two forward passes through the network. Due to the inherent randomness of Dropout, the neural activation states differ between the two passes, resulting in two distinct synergy probability prediction distributions, *p*^(1)^ and *p*^(2)^. The final composite loss function is defined as the weighted sum of the Binary Cross-Entropy (BCE) loss and the symmetric Kullback-Leibler (KL) divergence loss:

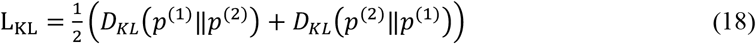

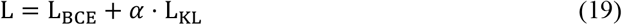

Where

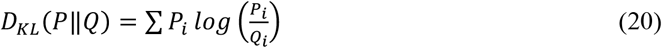

denotes the standard KL divergence, *L*_*BCE*_ is the average cross-entropy loss of the two forward passes, and α is a hyperparameter controlling the intensity of the consistency regularization. By employing this approach, the model is not only compelled to fit the ground-truth synergy labels but also effectively restricts the prediction variance between sub-models. This significantly mitigates the overfitting issues frequently encountered in deep Graph Neural Networks and Transformer modules.
2. Model Training and Hyperparameter Optimization: We employed the AdamW optimizer for end-to-end training of the model, complemented by the OneCycleLR learning rate scheduler to accelerate convergence. To prevent overfitting, dropout was used as the primary regularization method and was applied extensively within the feature projection layers, between the graph attention layers of the MolecularGNN, and inside the DiffTransformerFusion module.

All experiments were conducted on an NVIDIA RTX 4090 GPU using 5-fold cross-validation. Within each fold, we employed an early stopping strategy to determine the optimal number of training epochs: if the accuracy (ACC) on the validation set did not improve for 25 consecutive epochs (patience=25), the training was terminated early, and the model weights from the best-performing epoch were loaded for final testing. The model’s key hyperparameters, such as learning rate, batch size, dropout rate, and structural parameters of each module (e.g., number of GAT heads, Transformer depth), were determined through systematic manual tuning and ablation studies to ensure optimal performance.

## 3. Experiments and Results

### 3.1. Experimental Setup

#### 3.1.1. Evaluation Metrics

To comprehensively evaluate model performance, we adopted seven metrics: Accuracy (ACC), Balanced Accuracy (BACC), F1-score, Matthews Correlation Coefficient (MCC), Area Under the ROC Curve (AUC), Cohen’s Kappa (Kappa), and Area Under the Precision-Recall Curve (AP).

#### 3.1.2. Baseline Models

DeepSynergy [8]: This model concatenates chemical features of a drug pair and genomic features of a cell line into a single input vector, which is then processed by a feed-forward neural network to predict the synergy score.

MRGNN [20]: This model combines a multi-resolution graph neural network with dual LSTMs to simultaneously learn multi-scale drug features and their pairwise interactions.

EPGCN-DS [21]: This model uses an expressive graph convolutional network to learn atom-level features, which are then summed and processed by a Deep Sets decoder for interaction prediction.

DeepDDS [9]: This model concatenates drug embeddings from a graph attention network with cell line embeddings from an MLP for final synergy prediction.

MatchMaker [22]: This model separately processes each drug’s chemical features alongside cell line gene expression profiles in parallel subnetworks before merging their outputs for synergy prediction.

RWRDC [23]: This model constructs a drug network using only known synergistic combinations and then employs a Random Walk with Restart algorithm to score potential pairs based on their topological proximity within the network.

DFFNDDS [10]: This model employs a dual-feature fusion network, using parallel multi-head attention and highway modules, to process combined drug and cell line features for synergy prediction.

#### 3.1.3 Implementation Details

To ensure a fair comparison, all baseline models were reproduced by strictly following the hyperparameter settings described in their original papers or were carefully tuned on our validation set. For models originally designed for regression tasks, we modified their final layer to a Sigmoid function and used a binary cross-entropy (BCE) loss to adapt them to our binary classification setting. All models were implemented using PyTorch and PyTorch Geometric. The processing of drug molecules, including SMILES parsing, molecular graph construction, and fingerprint generation, was performed using the RDKit library, while the calculation of topological features relied on the NetworkX library. SMILES sequence encoding was handled using the Sentence-Transformers and Transformers libraries. The experiments were conducted on an NVIDIA RTX 4090 GPU. To ensure reproducibility, we fixed all random seeds to 42.

### 3.2. Overall Performance Comparison

We evaluated TopoFuseNet and all baseline models on two datasets: DrugCombDB and DrugComb. As shown in Table 1, TopoFuseNet significantly outperforms all baseline models on the DrugCombDB dataset. Paired t-tests confirm its superiority over the strongest baseline, DFFNDDS, across all seven reported metrics. This advantage is particularly pronounced in key metrics for imbalanced data, with AUPRC improving by 1.3% (p < 0.001) and the F1-score increasing by 2.4% (p < 0.05). This indicates that our method’s integration of multi-scale topological structures yields more discriminative drug representations. Similar performance advantages are observed on the DrugComb dataset (results are presented in Table S1 of Supplementary Material).

**Table 1.** Performance comparison of TopoFuseNet and baseline models on the DrugCombDB dataset with random splitting.

| Methods | ACC | BACC | F1 | AUC | MCC | Kappa | AP |
| --- | --- | --- | --- | --- | --- | --- | --- |
| TopoFuseNet | **0.869 (0.005)\*** | **0.833 (0.005)\*** | **0.771 (0.006)\*** | **0.925 (0.004)\*** | **0.680 (0.010)\*** | **0.680 (0.010)\*** | **0.861 (0.005)\*** |
| DFFNDDS | 0.862 (0.002) | 0.818 (0.003) | 0.753 (0.004) | 0.917 (0.001) | 0.660 (0.003) | 0.658 (0.003) | 0.850 (0.003) |
| DeepDDS | 0.819 (0.004) | 0.770 (0.005) | 0.681 (0.004) | 0.870 (0.004) | 0.557 (0.007) | 0.555 (0.007) | 0.775 (0.004) |
| DeepSynergy | 0.753 (0.003) | 0.627 (0.003) | 0.432 (0.007) | 0.737 (0.003) | 0.336 (0.005) | 0.300 (0.006) | 0.578 (0.004) |
| MRGNN | 0.804 (0.005) | 0.732 (0.007) | 0.627 (0.011) | 0.839 (0.005) | 0.506 (0.010) | 0.497 (0.011) | 0.721 (0.004) |
| EPGCNDS | 0.795 (0.005) | 0.719 (0.009) | 0.606 (0.013) | 0.826 (0.004) | 0.481 (0.014) | 0.472 (0.016) | 0.700 (0.008) |
| Matchmaker | 0.833 (0.004) | 0.779 (0.005) | 0.697 (0.006) | 0.882 (0.002) | 0.587 (0.007) | 0.584 (0.008) | 0.798 (0.002) |
| RWRDC | 0.474(0.001) | 0.605(0.003) | 0.512(0.005) | 0.607(0.005) | 0.233(0.010) | 0.143(0.004) | 0.393(0.004) |

Asterisks indicate statistical significance compared to DFFNDDS (paired t-test). \*: $p < 0.05$

Beyond the random dataset split, we further validated the model’s robustness under the four aforementioned cold-start settings with progressive difficulty. The experimental results reveal TopoFuseNet’s performance gradient when tackling tasks of varying cognitive difficulty. Figure 2 illustrates the core metric trends of our model alongside three representative baseline models for clear visual comparison (comprehensive results encompassing all 8 baseline models and 7 evaluation metrics are detailed in Tables S2-S5 of Supplementary Material):

1. Basic Combination Prediction (Generalized Combination Cold Start): This setting solely requires that the test combinations are absent from the training set. Under these conditions, TopoFuseNet achieves an AUC of 0.882. Because no strict limitations are placed on individual drugs during splitting, unknown individual drugs may occasionally be introduced into the test set. Despite this interference, the model maintains a high level of prediction accuracy.
2. Novel Combination Prediction with a Complete Vocabulary (Global Warm Start): To eliminate the interference of new drugs, this setting mandates that all individual drugs within the dataset have appeared in the training set (i.e., the model recognizes all drugs). In this “drug repurposing” scenario, the model yields an AUC of 0.881. This demonstrates that when the fundamental elements of the entire chemical space are known, the model can effectively bridge unknown interaction connections.
3. Interactive Logic Reasoning Scenario (Strict Combination Cold Start): This scenario evaluates pure reasoning under stringent conditions. Unlike the previous two, this setting not only necessitates entirely new test combinations but also strictly filters the test set, mandating that every individual drug in a test pair must have appeared in other pairings within the training set. This rigorous condition completely strips away the randomness of “learning new drug features,” purely evaluating the model’s capacity to reason about interactions between known drugs. Remarkably, TopoFuseNet’s AUC remains stable at 0.882 under these conditions. This proves that the model’s high scores are by no means reliant on data leakage or rote memorization; rather, it has genuinely learned the matching rules between molecular structures via the MolecularGNN module. As long as the individual drugs are known, the model can accurately deduce their pairing outcomes.
4. Structural Generalization Scenario (Drug-Level Cold Start): In the most challenging drug-level cold start setting, the model faces completely unknown chemical entities, demanding a strong inductive capability from structure to function. Although all baseline models experience an expected performance drop under this setting, TopoFuseNet still exhibits significant resilience ability, sustaining an AUC of 0.734 and outperforming the runner-up model, DFFNDDS (0.727). This relative advantage validates that, thanks to the integration of multi-scale topological features, the model has indeed learned universal chemical interaction principles. This highlights the model’s practical application value in screening potential synergistic drugs for completely novel (First-in-Class) molecules.

**Figure 2.**
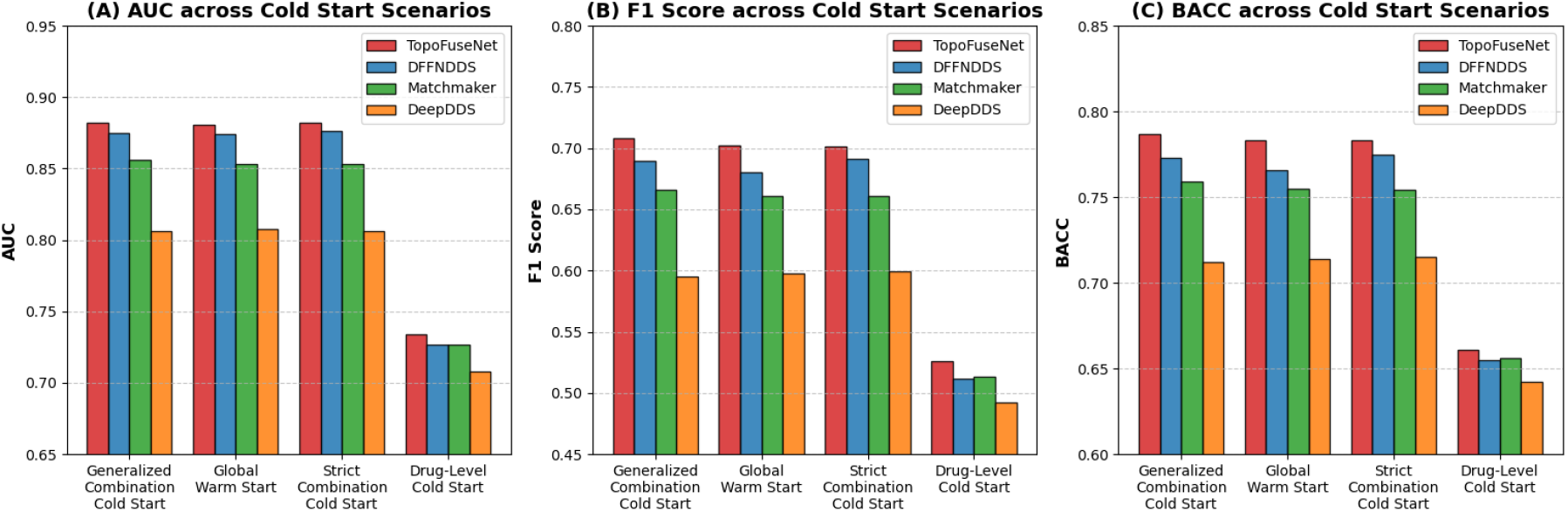
Comparison of core metrics between TopoFuseNet and three representative baseline models across four cold-start scenarios.

### 3.3. Ablation Studies

The stable predictive performance demonstrated across various data splitting scenarios prompted us to further investigate the specific contributions of the internal modules within TopoFuseNet. To validate the necessity and contribution of each proposed innovative module, we conducted a series of ablation studies. As shown in Table 2, removing or replacing any core module results in varying degrees of performance degradation, which strongly corroborates the rationality and effectiveness of our design.

1. Multimodal denoising advantage of the Differential Transformer: Replacing the Differential Transformer with a standard Transformer leads to a 1.0% drop in the F1-score. Compared to the standard Transformer, the differential mechanism effectively cancels out common-mode noise during multimodal feature fusion by configuring paired primary and auxiliary attention heads. Specifically, the model obtains the final attention weights by computing the difference between two attention matrices:

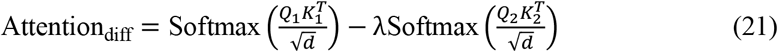 In our model, GNN structural embeddings, fine-tuned BERT semantic sequences, and traditional molecular fingerprints are aggregated. Although these heterogeneous data are complementary, information redundancy is inevitable. Traditional Transformers are prone to attention dispersion amidst such common-mode noise. In contrast, the Differential Transformer allows the primary attention head to capture global features, while the auxiliary attention head specifically fits and cancels out the redundant noise, thereby achieving a sparser and purer cross-modal drug representation.
2. Chemical intuition behind Group Centrality-guided hierarchical pooling: Replacing the Group Centrality-based pooling with standard average pooling causes a 0.4% decrease in the F1-score, demonstrating the effectiveness of the hierarchical pooling strategy that focuses on functional groups using group centrality. Our model employs Group Degree Centrality, Group Closeness Centrality, and Group Betweenness Centrality to quantify the topological importance of chemical groups. At the physicochemical level, Group Degree Centrality effectively reflects the reaction accessibility and surface exposure of drug groups; highly exposed groups are more likely to become the pharmacophoric core for intermolecular interactions. Group Closeness Centrality reveals the potential of a group in transmitting electronic effects (e.g., conjugation and inductive effects); groups located in central positions (e.g., core aromatic rings) can more efficiently regulate the overall electron cloud distribution. Group Betweenness Centrality helps identify groups that act as the molecular skeleton (e.g., long-chain alkanes or spiro rings); although these groups may not directly participate in bond formation, they dictate the spatial conformation and hydrophobic interactions of the molecule. In summary, traditional average pooling obscures the locally significant signals of key pharmacophores, whereas Group Centrality pooling endows the model with the ability to weight and perceive molecular groups based on graph-theoretic features, thereby learning more discriminative molecular representations.
3. Structural guidance of topology-aware attention: Removing the topology-aware attention module also leads to performance degradation, confirming the value of explicitly injecting topological priors into the GNN. This model adopts “Second Order Centrality” as a bias term during the GAT message-passing process. In a molecular graph, a lower Second Order Centrality value indicates that the atom is more inclined to constitute a stable molecular skeleton; conversely, a higher value signifies that the atom is located at the topological periphery, making it more likely to belong to an active reactive side chain or a modifying group. The introduction of this topological bias enables the GAT to a priori perceive the structural differences between the core molecular skeleton and peripheral active sites at the early stage of aggregation. Evidence shows that this distinction effectively boosts model performance.
4. Necessity of multimodal feature fusion: Removing the ECFP molecular fingerprints and the fine-tuned BERT embeddings results in F1-score drops of 0.6% and 0.4%, respectively. This indicates that although the highly customized MolecularGNN has already extracted rich graph topological information, substructure-based fingerprint matching (ECFP) and sequence-syntax-based pre-trained language model representations (BERT) still provide irreplaceable complementary perspectives, corroborating the necessity of employing a multimodal feature fusion architecture in this model.

**Table 2.** Ablation studies of TopoFuseNet’s core modules.

| Methods | ACC | BACC | F1 | AUC | MCC | Kappa | Ap |
| --- | --- | --- | --- | --- | --- | --- | --- |
| TopoFuseNet | **0.869(0.005)** | **0.833(0.005)** | **0.771(0.006)** | **0.925(0.004)** | **0.680(0.010)** | **0.679(0.010)** | **0.861(0.005)** |
| w/o Differential Transformer | 0.865(0.003)* | 0.827(0.002)* | 0.763(0.002)* | 0.923(0.002)* | 0.671(0.004)* | 0.700(0.003)* | 0.859(0.002)* |
| w/o Group Centrality Pooling | 0.867(0.005) | 0.830(0.008)* | 0.768(0.009)* | 0.924(0.003)* | 0.676(0.012) | 0.675(0.013) | 0.859(0.004)* |
| w/o Topology-Aware Attention | 0.868(0.003) | 0.830(0.003) | 0.768(0.003)* | 0.924(0.002) | 0.677(0.005)* | 0.676(0.006)* | 0.858(0.001)* |
| w/o Molecular Fingerprint (FP) | 0.867(0.004)* | 0.829(0.005) | 0.766(0.005) | 0.925(0.003) | 0.675(0.007)* | 0.673(0.008)* | 0.862(0.003) |
| w/o Fine-tuned BERT | 0.867(0.004) | 0.831(0.003) | 0.768(0.004) | 0.924(0.003)* | 0.676(0.007) | 0.675(0.007) | 0.859(0.004)* |

Asterisks indicate statistical significance compared to TopoFuseNet (paired t-test). \*: $p < 0.05$

### 3.4. Parameter Sensitivity Analysis

After confirming the necessity of the core modules, to evaluate the robustness of TopoFuseNet and determine the optimal hyperparameter configuration, we conducted a sensitivity analysis on two key structural parameters: the feature projection dimension (*d*_*proj*_) prior to the Differential Transformer and the depth of the Transformer module (*L*). The relevant comparative experiments were conducted on the DrugCombDB dataset, and the performance trends are illustrated in Figure 3.

**Figure 3.**
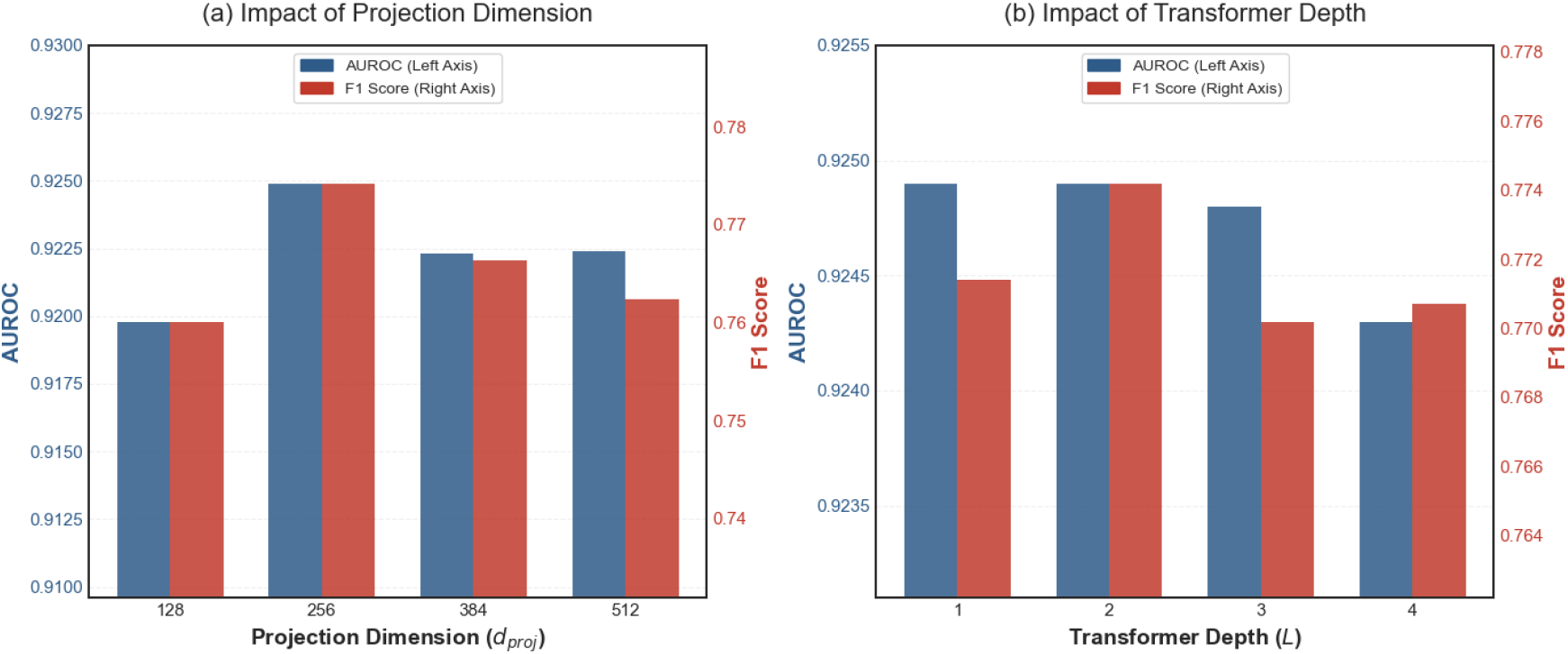
Hyperparameter sensitivity analysis. (a) Impact of feature projection dimension on model performance; (b) Impact of Differential Transformer depth on model performance.

First, we investigated the impact of the projection dimension *d*_*proj*_ ∈ {128,256,384,512} on model performance. As shown in Figure 3(a), the model performance exhibits an initial increase followed by a decrease as the dimension grows, peaking distinctly at *d*_*proj*_ = 256. When the feature dimension is relatively low (e.g., *d*_*proj*_ = 128), the limited feature space restricts the representational capacity of the model, resulting in an F1-score of only 0.7601 (compared to 0.7742 for the optimal configuration). However, indiscriminately increasing the projection dimension does not yield continuous performance gains; when *d*_*proj*_ increases to 512, the F1-score drops back to 0.7624. This phenomenon indicates that a 256-dimensional feature space can adequately encode complex chemical and biological semantics while effectively circumventing the risk of overfitting associated with high-dimensional feature spaces.

Second, we evaluated the impact of the Differential Transformer depth (*L* ∈ {1, 2, 3, 4}). The results in Figure 3(b) confirm that a depth of *L* = 2 achieves the optimal predictive performance. Specifically, compared to a single-layer network (*L* = 1), the introduction of a second Transformer layer significantly enhances the model’s ability to capture complex cross-modal interactions between drugs and cell lines, elevating the F1-score to its peak of 0.7742. However, as the network deepens further (*L* ≥ 3), the model performance experiences a slight degradation (e.g., the F1-score drops to approximately 0.770). This performance deterioration is highly consistent with the “oversmoothing” phenomenon commonly observed in graph representation learning and deep attention mechanisms, where excessive message passing and fusion inadvertently dilute feature distinctiveness without providing additional informative gains. Balancing predictive accuracy against computational overhead, we ultimately established *L* = 2 as the default configuration for the model, ensuring high accuracy while maintaining operational efficiency.

### 3.5. Visualization of Model Representations and Biological Sensitivity Analysis

The aforementioned quantitative experiments have fully validated the superiority of the model architecture. To go beyond mere performance metrics and visually evaluate the quality of the multimodal representations learned by TopoFuseNet, as well as its sensitivity to biological contexts, we conducted an in-depth visual analysis of the test set samples.

1. Discriminative Power in Latent Space: We extracted the penultimate layer feature vectors prior to the final classification layer (i.e., the multimodal embeddings fused by the Differential Transformer) and projected them into a two-dimensional space using the t-SNE algorithm. As shown in Figure 4(A), the sample points are colored according to their predicted synergistic probabilities. It can be clearly observed that the latent manifold learned by the model exhibits a distinct clustering structure driven by the predicted probabilities: samples with high synergistic probabilities (yellow regions) and antagonistic/non-synergistic samples (dark purple regions) are effectively separated in the feature space. This high degree of feature discriminability indicates that, by integrating multi-scale topological and sequential information, TopoFuseNet successfully maps the original complex inputs into highly discriminative feature representations, laying a solid foundation for accurate final classification.
2. Tissue-Specific Synergy Patterns: Drug synergy depends not only on the chemical structures of the drug molecules but also heavily on the biological context of the cell lines. To verify whether the model captures this context dependency, we further analyzed the distribution of predicted scores and the predicted synergy rates across different tissue types.

**Figure 4.**
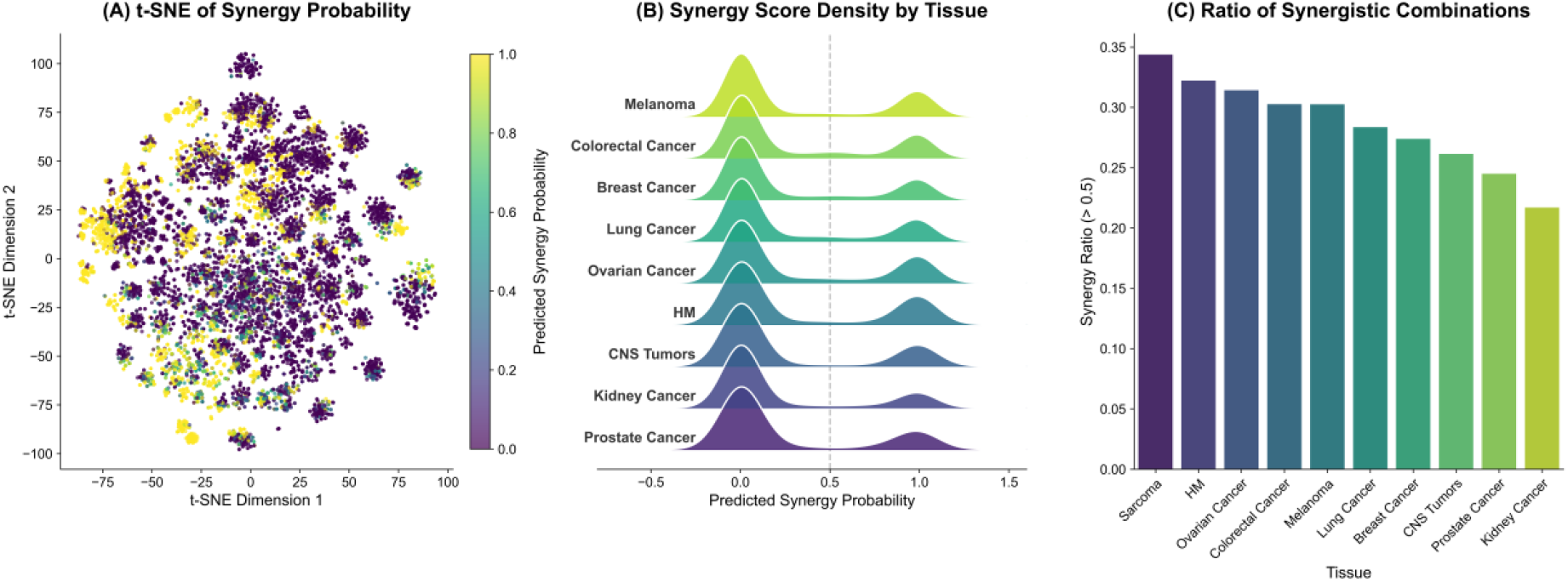
Visualization of the latent feature space and tissue-specific synergy patterns. (A) t-SNE projection of the penultimate layer’s multimodal embedding vectors, colored by predicted synergy probability. (B) Ridge plots displaying the density distribution of predicted synergy scores across different cancer tissue types. The dashed gray line represents the classification threshold of 0.5. (C) Proportion of predicted synergistic combinations (probability > 0.5) ranked by tissue type.

As illustrated in Figure 4(B), although the ridge plots of predicted score densities for different cancer tissue types all exhibit a bimodal distribution concentrated at both extremes—reflecting the model’s stable high confidence in cross-tissue predictions—they demonstrate significant biological heterogeneity in their specific synergistic tendencies. Further quantitative analysis (Figure 4(C)) reveals the differences in synergistic response rates among various cancer types:

Sarcoma (synergy rate ∼34.4%) and Hematologic Malignancies (synergy rate ∼32.2%) exhibit the highest predicted synergistic proportions, whereas Kidney Cancer (synergy rate ∼21.7%) is relatively low. This predictive trend aligns with clinical observations where certain tumor types are more sensitive to combination chemotherapy. This result strongly demonstrates that TopoFuseNet does not merely memorize drug structures, but effectively integrates cell line gene expression features, thereby enabling personalized synergy predictions tailored to specific biological contexts.

### 3.6. Case Studies: Multi-Scale Interpretability

The clustering distribution in the feature space initially validates the effectiveness of the model’s representations. To further investigate how TopoFuseNet conducts biological reasoning at both the microscopic atomic scale and the macroscopic modal scale, we performed a comprehensive interpretability analysis on three distinct drug combinations encompassing various cancer types and synergy mechanisms. By visualizing the microscopic atomic-level attention weights of the GNN module and the macroscopic feature fusion patterns of the Differential Transformer, we demonstrate how the model hierarchically deciphers complex pharmacological interactions.

#### 3.6.1. Microscopic Level

Deciphering Key Pharmacophores via the GNN Attention Mechanism At the atomic level, the attention weights learned by our topology-enhanced GAT module exhibit exceptionally high consistency with known Structure-Activity Relationships (SAR) and drug resistance mechanisms (Figure 5).

**Figure 5.**
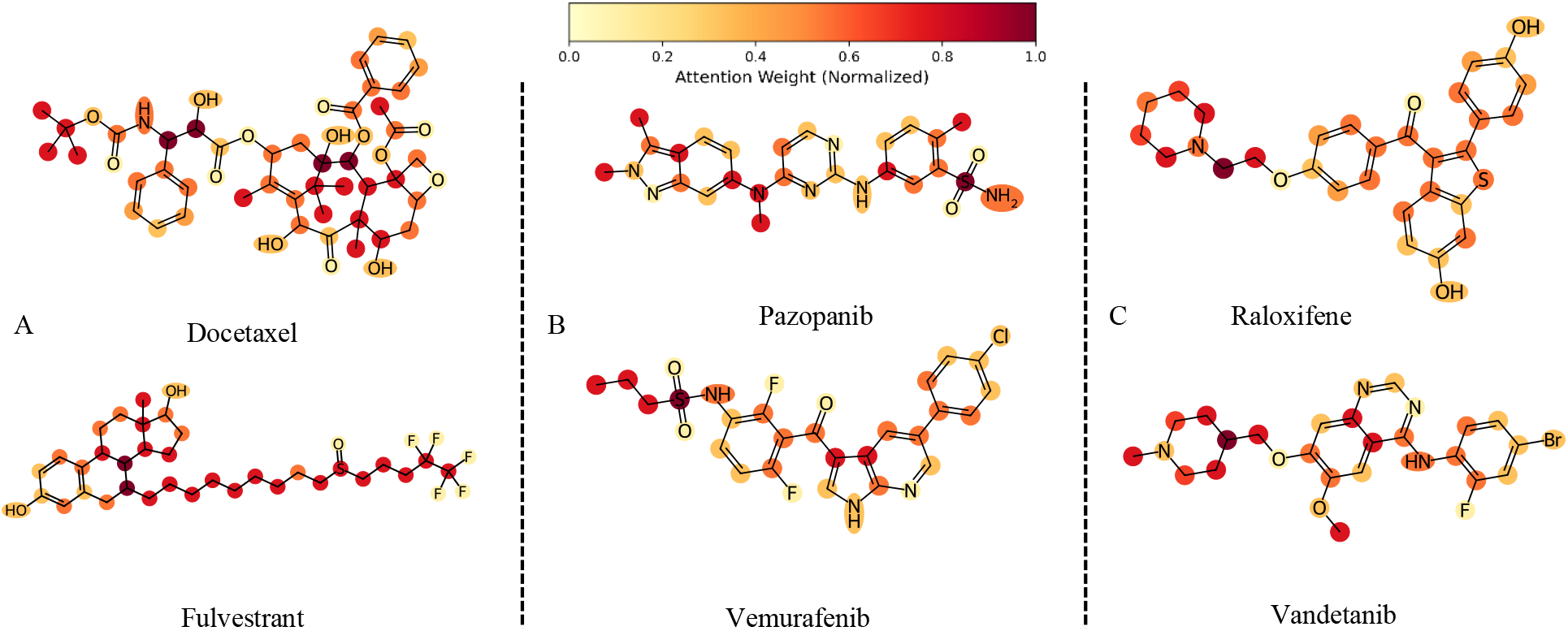
Visualization of molecular atomic attention weights for three synergistic drug combinations. (A) Docetaxel + Fulvestrant; (B) Pazopanib + Vemurafenib; (C) Raloxifene + Vandetanib.

Case 1: Overcoming Drug Resistance (Docetaxel + Fulvestrant in U251 cells). In the context of P-gp overexpressing glioblastoma cells[24], the model exhibits remarkable biological specificity. For fulvestrant (Figure 5A), the model pays little attention to the steroid core responsible for estrogen receptor (ER) binding [25] (which is less relevant in the current context), but instead assigns the maximum attention weight (1.00) to the alkyl sulfinyl side chain. This accurately reflects the pharmacological reality: this hydrophobic chain competitively blocks the P-gp efflux pump, thereby reversing drug resistance[26]. Meanwhile, for docetaxel (Figure 5A), the model highlights the C13 ester side chain, which is a critical group for microtubule stabilization[27]. Consequently, the model infers a “pharmacokinetic sensitization” mechanism: fulvestrant blocks the efflux pump, enabling the cytotoxic docetaxel to accumulate and exert its efficacy[28].

Case 2: Paradoxical Activation Blockade (Vemurafenib + Pazopanib in the SK-MEL-2 cell line). In NRAS-mutant melanoma, single BRAF inhibition induces paradoxical activation[29], whereas the model identifies a structural solution. For pazopanib (Figure 5B), the model pays high attention to the sulfonamide moiety and the indazole-methyl group (weights 0.75-1.00), which are key features for Type II kinase binding that lock BRAF in the inactive DFG-out conformation[30, 31]. For vemurafenib (Figure 5B), the model correctly prioritizes the propyl side chain (weight 0.75), which is crucial for its selectivity towards the BRAF V600E pocket[32]. This implies a “pan-RAF blockade” strategy: pazopanib prevents the dimerization-induced transactivation typically caused by vemurafenib in this specific genetic context[33, 34].

Case 3: Non-canonical Synergy (Raloxifene + Vandetanib in the HS 578T cell line). In ER-negative triple-negative breast cancer (TNBC) cells, the model downplays raloxifene’s classical hormonal mechanism, assigning a lower weight to the phenolic hydroxyl group responsible for ER binding[35]. Conversely, it focuses heavily on the piperidine-ethoxy side chain (weight 1.00), a known P-gp/BCRP inhibitor[36]. Concurrently, for vandetanib, the model focuses on the N-methylpiperidinyl-methoxy side chain (weight 1.00), which is the primary recognition site for the efflux pump[37]. This reveals a complex “side chain-side chain” interaction logic, wherein raloxifene protects vandetanib from efflux [38], thereby achieving intracellular kinase inhibition[39].

#### 3.6.2. Macroscopic Level: Orchestrating Multimodal Reasoning via Differential Attention

Beyond atomic features, the attention heatmaps from the Differential Transformer (Figure 6) reveal that TopoFuseNet employs a dynamic and context-aware reasoning strategy. Depending on the biological context and drug properties, the model shifts its focus across different modalities, exhibiting three distinct reasoning patterns.

1. Pattern 1: Contextual Feature Filtering. As shown in Figure 6A, for the DNA damage repair combination, the model demonstrates a noise suppression mechanism. Notably, the explicit sequence information of docetaxel (SMILES_1) is actively suppressed by the fingerprint feature (FP_1) (with a negative attention value of -0.07). This indicates that for complex macromolecules like taxanes, the model filters out noisy sequence data. In contrast, the GNN representations (GNN_1, GNN_2) maintain a stable and positive attention distribution (0.06-0.10) across all attention heads. This suggests that hierarchical graph features provide a consistent topological baseline, keeping the prediction results stable amidst the fluctuations of explicit 1D features.
2. Pattern 2: Context-Guided Modality Prioritization. For the kinase inhibitor combination in melanoma (Figure 6B), the attention pattern highlights a strong dependence on the biological context. The cell line context exclusively allocates its maximum attention (0.14) to the sequence of pazopanib (SMILES_2) rather than vemurafenib. This aligns with the biological reality: in NRAS-mutant SK-MEL-2 cells, intrinsic resistance to the BRAF inhibitor (vemurafenib) shifts the therapeutic burden onto the second drug[29]. The model accurately discerns that, within this specific genetic context, the efficacy of the combination therapy critically depends on the multi-target kinase inhibitor, pazopanib.
3. Pattern 3: Graph-Level Structural Complementarity. In the complex scenario of triple-negative breast cancer (TNBC) (Figure 6C), due to the lack of classical target binding mechanisms (e.g., raloxifene’s modulation of ER)[35], the model shifts its reliance toward higher-order structural features. We observe the strongest GNN-GNN interaction among the three cases, where GNN_2 (vandetanib) exhibits intense attention to itself (0.10) and to GNN_1 (raloxifene) (0.08). This enhanced interdependence between graph modalities indicates that, in the absence of explicit targets (like ER), the model pivots to higher-order topological matching to identify spatial complementarity between molecules.

**Figure 6.**
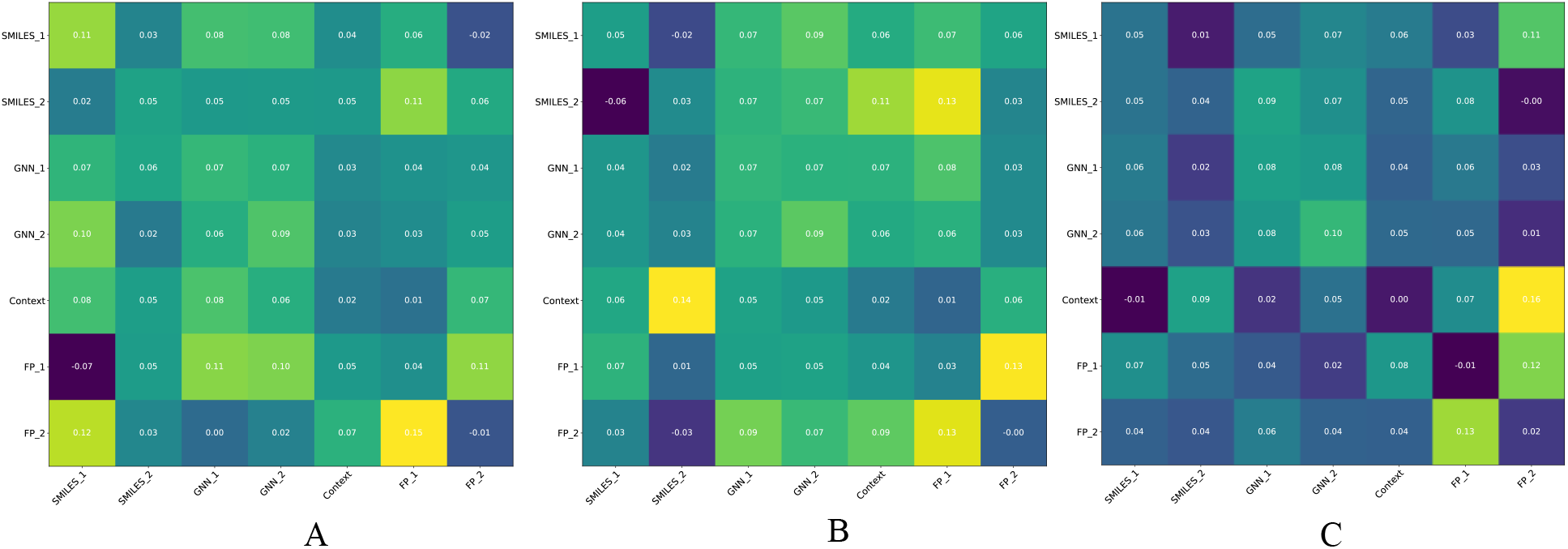
Differential Transformer feature fusion attention heatmaps for three synergistic drug combinations. (A) Docetaxel + Fulvestrant; (B) Pazopanib + Vemurafenib; (C) Raloxifene + Vandetanib.

In summary, TopoFuseNet does more than merely memorize chemical patterns; it functions as an interpretable chemical reasoning engine. It autonomously identifies key pharmacophores at the microscopic level and dynamically modulates cross-modal attention at the macroscopic level—filtering sequence redundancy, prioritizing context-relevant features, and evaluating global graph complementarity. This multi-scale interpretability tightly couples the model’s computational attention with solid pharmacological mechanisms, significantly enhancing the reliability of deep learning in drug combination discovery.

### 3.7. Zero-Shot Cross-Domain Generalization and Virtual Screening Capability Analysis

The preceding multi-scale interpretability analysis demonstrated that the model can effectively capture key pharmacophores and dynamically orchestrate multimodal information. To further evaluate whether TopoFuseNet has learned generalizable underlying principles of molecular interactions that transcend specific cell line contexts, we conducted a zero-shot cross-domain transfer experiment on the Ryu (TWOSIDES) clinical drug-drug interaction (DDI) dataset (191,388 positive pairs and an equal number of random negative samples). To construct a rigorous out-of-distribution (OOD) scenario, we masked the specific cell line features (replacing them with a global mean vector *C*_*mean*_), forcing the model to rely solely on the multi-scale topological and sequential features of the molecules for pure chemical structure reasoning.

Although the model’s global AUC is constrained (0.5790) due to the semantic misalignment of labels between “anti-cancer synergy” and “generalized side effects (including antagonism)”, TopoFuseNet still exhibits exceptional discriminative power. As shown in Figure 7, the predicted probability density of true interacting pairs (red line) exhibits a significant shift toward the high-score region on the right compared to random negative samples (blue line). This visual observation is rigorously confirmed by the Kolmogorov-Smirnov test (*KS* = 0.1215, *p* < 0.001), and the mean predicted probability score of positive samples is significantly higher (0.3998 vs. 0.3381), indicating that the model successfully captures the structural prerequisites for intermolecular interactions.

**Figure 7.**
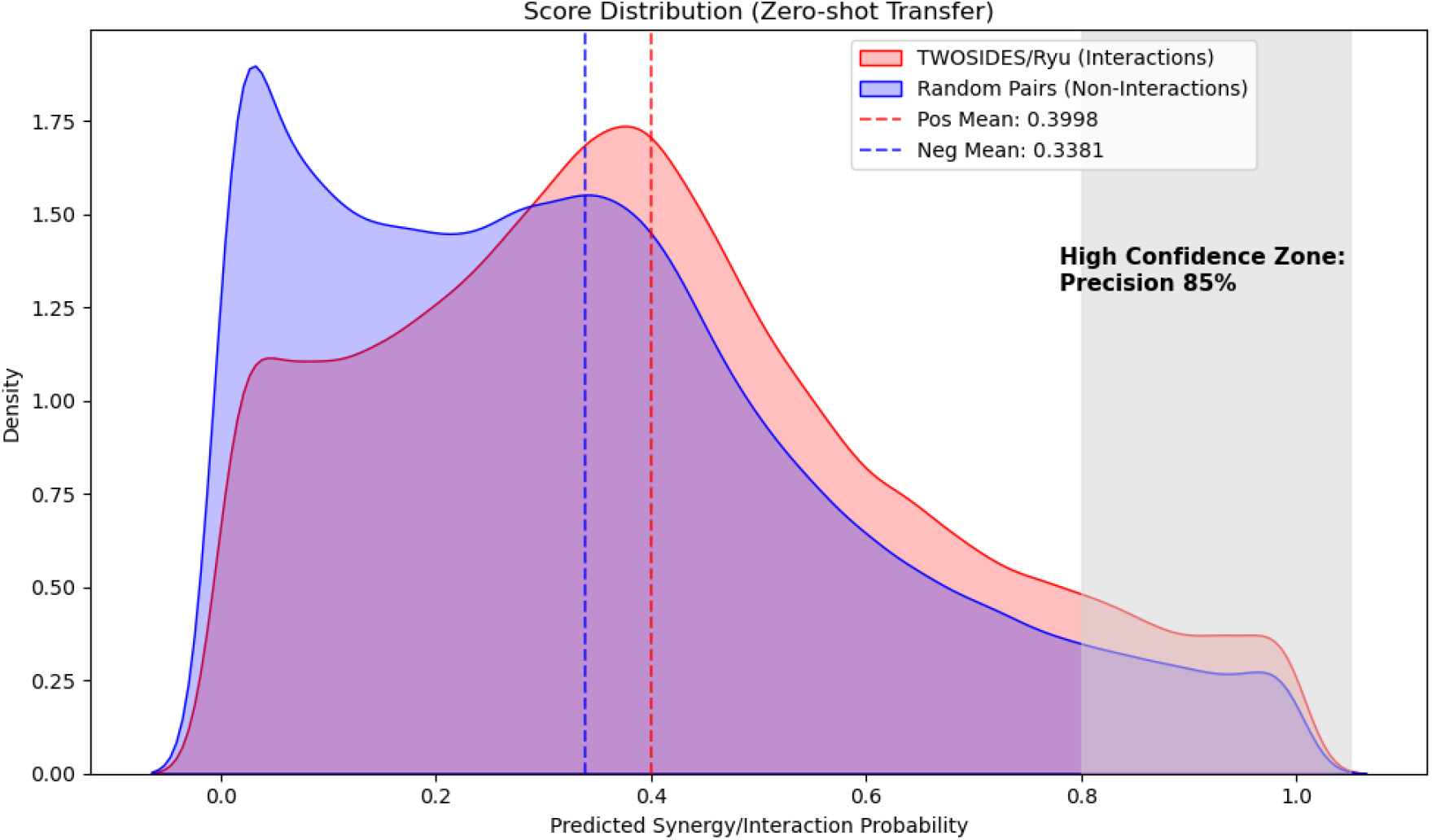
Predicted probability density distribution in the zero-shot cross-domain transfer task. (The red area represents true drug interaction pairs, while the blue area represents randomly generated negative sample combinations.)

More importantly, in the Early Recognition scenario—which is highly valued in practical drug discovery—the model demonstrates its immense potential as an efficient virtual screening engine. Within the model’s extremely high confidence interval, the precision of the Top-100 predictions reaches an impressive 85.00%, and the Top-500 maintains a precision of 71.20%. Furthermore, the model achieves an Enrichment Factor of 1.17 in the Top 1% prediction interval, meaning its efficiency in mining strong interaction signals from massive background noise is significantly superior to random screening. These results robustly confirm that TopoFuseNet has successfully encoded universal physicochemical rules, such as spatial pharmacophore matching, and possesses practical application value in providing high-confidence candidate rankings for the exploration of novel clinical combination therapies.

### 3.8. Exploration of Practical Application Scenarios: Discovery of Novel Anti-Cancer Drug Combinations

The results of the zero-shot cross-domain transfer experiment preliminarily validated the potential of TopoFuseNet as a virtual screening tool. Building upon this finding, to further evaluate the practical application potential of TopoFuseNet in guiding preclinical drug screening and discovering novel combination therapies, we designed a large-scale in silico screening experiment. Going beyond the test set, we aimed to explore whether the model could identify potentially synergistic drug combinations from a massive, unexplored space.

We collected all 1,084 unique single drugs from the DrugCombDB dataset. Through pairwise pairing, we constructed a massive candidate library comprising 579,930 potential drug combinations. To ensure the rigor of the evaluation, we strictly excluded the drug pairs already existing in the original DrugCombDB dataset, ensuring that all input data was entirely novel to TopoFuseNet (Novel Combinations).

We evaluated these novel drug pairs in the context of nine representative cell lines across three high-incidence cancers for prediction. These nine cell lines encompass a broad range of genetic backgrounds, including:

Lung Cancer: A549, NCI-H460, NCI-H2122

Colorectal Cancer: HT-29, KM12, DLD-1 Breast Cancer: BT-549, T-47D, MDA-MB-468

For each cell line, we ranked all candidate combinations based on the predicted probabilities output by TopoFuseNet and selected the top 3 high-confidence synergistic combinations for detailed literature validation.

The results are highly encouraging. Through extensive searches of PubMed and clinical trial databases, we found that most of the Top-3 combinations predicted by the model are supported by direct or indirect evidence in existing biomedical literature. For a complete list of all evaluated novel combinations and their supporting literature evidence, please refer to Supplementary Tables S6-S15. Table 3 presents some representative novel synergistic combinations predicted by the model alongside their supporting literature evidence. This result robustly demonstrates that TopoFuseNet not only fits the training data but, more importantly, successfully captures the complex non-linear interactions between molecular structures (especially the topological features of key functional groups) and cell line genomic features. This empowers it to discover potentially highly efficacious combination therapies in uncharted spaces.

**Table 3.** Representative novel synergistic drug combinations predicted by TopoFuseNet and their supporting literature evidence.

| Cancer Type | Cell Line | Drug A | Drug B | Evidence |
| --- | --- | --- | --- | --- |
| Lung Cancer | NCI-H460 | etoposide | melphalan | In P388 leukemia cells, the combination of etoposide and melphalan has been confirmed to exhibit strong synergism via median-effect analysis. In patients with neuroblastoma, high-dose chemotherapy regimens incorporating these two drugs (e.g., the CEM regimen) have demonstrated superior clinical response rates[40, 41]. |
| Colorectal Cancer | HT-29 | Mitoxantrone | Chloroquine | Mitoxantrone induces protective autophagy. Chloroquine blocks the autophagic process and sensitizes HT-29 cells to chemotherapy and radiotherapy. Direct studies demonstrate that the combination of chloroquine and mitoxantrone exerts a profound synergistic effect (CI = 0.40) in drug-resistant models[42, 43]. |
| Breast Cancer | T-47D | Vemurafenib | Tideglusib | Literature confirms that the inhibition of GSK-3 (via either siRNA or the pharmacological inhibitor AR-A014418) can sensitize melanoma cells to Vemurafenib/Sorafenib and enhance apoptosis. Tideglusib, a similar GSK-3 inhibitor, has demonstrated single-agent activity and chemosensitizing effects in neuroblastoma[44, 45]. |

## 4. Discussion

This study successfully developed a novel deep learning framework named TopoFuseNet for the precise prediction of drug synergy. Experimental results demonstrate that by systematically integrating multi- scale topological information, our model significantly outperforms existing methods in terms of performance.

The core reason for our model’s superior performance lies in overcoming the limitation of traditional GNNs, which conventionally treat molecules as flat graphs. By introducing group centrality to model functional groups and designing a multi-pathway fusion strategy, TopoFuseNet is capable of learning hierarchical drug representations that more closely align with their chemical nature. Ablation studies and parameter sensitivity analyses clearly demonstrate that each proposed innovative module, particularly the entire topological fusion paradigm, is crucial for performance enhancement. Case studies further reveal that the model is not only accurate in its predictions, but its internal attention mechanisms also align with known pharmacological knowledge, providing robust evidence for the model’s interpretability. More importantly, the zero-shot cross-domain transfer and the in silico screening experiment at a scale of nearly 580,000 fully substantiate its immense potential in real-world preclinical drug discovery.

Despite the outstanding performance of TopoFuseNet, some limitations remain. First, the model’s performance relies on the quality and completeness of existing databases, and data noise and biases may affect the prediction results. Second, although we have preliminarily explored interpretability through case studies, the “black box” nature of deep learning models has not been fully resolved. Finally, this study primarily focuses on the binary classification of synergistic effects, without considering dose-effect relationships.

Future research can be pursued in the following directions: 1. Extending the model to multi-class classification (e.g., synergy, antagonism, additivity) and regression tasks (predicting synergy scores). 2. Integrating additional data modalities, such as proteomics data and clinical patient data, to construct a more comprehensive predictive model. 3. Applying the model to real-world drug screening pipelines and validating its predictive results through wet-lab experiments, ultimately establishing a computational-experimental feedback loop.

## 5. Conclusion

To address the limitation that existing drug synergy prediction methods fail to fully utilize hierarchical molecular structures and topological information, this study proposes a novel deep learning framework named TopoFuseNet. The core innovation of this model lies in introducing group centrality theory into chemoinformatics for the first time and designing a systematic, multi-pathway topological information fusion mechanism to learn more informative drug representations. Experiments on two large-scale benchmark datasets demonstrate that TopoFuseNet significantly outperforms existing state-of-the-art methods across multiple key metrics. Combined with its demonstrated multi-scale interpretability and zero-shot generalization capabilities, this work provides a more reliable computational tool for the large-scale, automated screening of synergistic drug combinations, holding the promise to promote the development of anti-cancer combination therapies and personalized medicine.

## Data Availability

The public datasets (DrugComb and DrugCombDB) analyzed during the current study are openly accessible. The source code of TopoFuseNet supporting the findings of this study is available from the corresponding author upon reasonable request.

## CRediT authorship contribution statement

Xiaoyang Shi: Methodology, Software, Validation, Formal analysis, Investigation, Data curation, Writing - original draft, Visualization. Qi Wang: Conceptualization, Supervision, Project administration, Funding acquisition, Writing - review & editing.

## References

[1] R. Bayat Mokhtari et al., “Combination therapy in combating cancer,” (in eng), Oncotarget, vol. 8, no. 23, pp. 38022–38043, Jun 6 2017, doi: 10.18632/oncotarget.16723.

[2] T.-C. Chou, “Drug Combination Studies and Their Synergy Quantification Using the Chou-Talalay Method,” Cancer Research, vol. 70, no. 2, pp. 440–446, 2010, doi: 10.1158/0008-5472.Can-09-1947.

[3] J. B. Fitzgerald, B. Schoeberl, U. B. Nielsen, and P. K. Sorger, “Systems biology and combination therapy in the quest for clinical efficacy,” Nature Chemical Biology, vol. 2, no. 9, pp. 458–466, 2006/09/01 2006, doi: 10.1038/nchembio817.

[4] D. Sun, W. Gao, H. Hu, and S. Zhou, “Why 90% of clinical drug development fails and how to improve it?,” Acta Pharmaceutica Sinica B, vol. 12, no. 7, pp. 3049–3062, 2022/07/01/ 2022, doi: 10.1016/j.apsb.2022.02.002.

[5] N. P. Tatonetti, P. P. Ye, R. Daneshjou, and R. B. Altman, “Data-Driven Prediction of Drug Effects and Interactions,” Science Translational Medicine, vol. 4, no. 125, pp. 125ra31–125ra31, 2012, doi: doi:10.1126/scitranslmed.3003377.

[6] F. Cheng and Z. Zhao, “Machine learning-based prediction of drug-drug interactions by integrating drug phenotypic, therapeutic, chemical, and genomic properties,” (in eng), J Am Med Inform Assoc, vol. 21, no. e2, pp. e278–86, Oct 2014, doi: 10.1136/amiajnl-2013-002512.

[7] A. Gottlieb, G. Y. Stein, Y. Oron, E. Ruppin, and R. Sharan, “INDI: a computational framework for inferring drug interactions and their associated recommendations,” (in eng), Mol Syst Biol, vol. 8, p. 592, Jul 17 2012, doi: 10.1038/msb.2012.26.

[8] K. Preuer, R. P. I. Lewis, S. Hochreiter, A. Bender, K. C. Bulusu, and G. Klambauer, “DeepSynergy: predicting anti-cancer drug synergy with Deep Learning,” (in eng), Bioinformatics, vol. 34, no. 9, pp. 1538–1546, May 1 2018, doi: 10.1093/bioinformatics/btx806.

[9] J. Wang, X. Liu, S. Shen, L. Deng, and H. Liu, “DeepDDS: deep graph neural network with attention mechanism to predict synergistic drug combinations,” (in eng), Brief Bioinform, vol. 23, no. 1, Jan 17 2022, doi: 10.1093/bib/bbab390.

[10] M. Xu et al., “DFFNDDS: prediction of synergistic drug combinations with dual feature fusion networks,” Journal of Cheminformatics, vol. 15, no. 1, p. 33, 2023/03/16 2023, doi: 10.1186/s13321-023-00690-3.

[11] J. Sun and H. Zheng, “HDN-DDI: a novel framework for predicting drug-drug interactions using hierarchical molecular graphs and enhanced dual-view representation learning,” BMC Bioinformatics, vol. 26, no. 1, p. 28, 2025/01/25 2025, doi: 10.1186/s12859-025-06052-0.

[12] M. Everett and S. Borgatti, “The Centrality of Groups and Classes,” Journal of Mathematical Sociology, vol. 23, pp. 181–201, 01/01 1999, doi: 10.1080/0022250X.1999.9990219.

[13] B. Zagidullin et al., “DrugComb: an integrative cancer drug combination data portal,” (in eng), Nucleic Acids Res, vol. 47, no. W1, pp. W43–w51, Jul 2 2019, doi: 10.1093/nar/gkz337.

[14] H. Liu, W. Zhang, B. Zou, J. Wang, Y. Deng, and L. Deng, “DrugCombDB: a comprehensive database of drug combinations toward the discovery of combinatorial therapy,” (in eng), Nucleic Acids Res, vol. 48, no. D1, pp. D871–d881, Jan 8 2020, doi: 10.1093/nar/gkz1007.

[15] J. Devlin, M.-W. Chang, K. Lee, and K. Toutanova, “BERT: Pre-training of Deep Bidirectional Transformers for Language Understanding,” Minneapolis, Minnesota, June 2019: Association for Computational Linguistics, in Proceedings of the 2019 Conference of the North American Chapter of the Association for Computational Linguistics: Human Language Technologies, Volume 1 (Long and Short Papers), pp. 4171–4186, doi: 10.18653/v1/N19-1423. [Online]. Available: https://aclanthology.org/N19-1423/ https://doi.org/10.18653/v1/N19-1423

[16] S. Chithrananda, G. Grand, and B. Ramsundar, “ChemBERTa: Large-Scale Self-Supervised Pretraining for Molecular Property Prediction,” p. 2010.09885 doi: 10.48550/arXiv.2010.09885.

[17] T. Gao, X. Yao, and D. Chen, “SimCSE: Simple Contrastive Learning of Sentence Embeddings,” Online and Punta Cana, Dominican Republic, November 2021: Association for Computational Linguistics, in Proceedings of the 2021 Conference on Empirical Methods in Natural Language Processing, pp. 6894–6910, doi: 10.18653/v1/2021.emnlp-main.552. [Online]. Available: https://aclanthology.org/2021.emnlp-main.552/ https://doi.org/10.18653/v1/2021.emnlp-main.552

[18] J. Barretina et al., “The Cancer Cell Line Encyclopedia enables predictive modelling of anticancer drug sensitivity,” (in eng), Nature, vol. 483, no. 7391, pp. 603–7, Mar 28 2012, doi: 10.1038/nature11003.

[19] T. Ye et al., “Differential Transformer,” p. 2410.05258 doi: 10.48550/arXiv.2410.05258.

[20] N. Xu, P. Wang, L. Chen, J. Tao, and J. Zhao, “MR-GNN: Multi-Resolution and Dual Graph Neural Network for Predicting Structured Entity Interactions,” p. 1905.09558 doi: 10.48550/arXiv.1905.09558.

[21] M. Sun, F. Wang, O. Elemento, and J. Zhou, “Structure-Based Drug-Drug Interaction Detection via Expressive Graph Convolutional Networks and Deep Sets (Student Abstract),” Proceedings of the AAAI Conference on Artificial Intelligence, vol. 34, no. 10, pp. 13927–13928, 04/03 2020, doi: 10.1609/aaai.v34i10.7236.

[22] H. I. Kuru, O. Tastan, and A. E. Cicek, “MatchMaker: A Deep Learning Framework for Drug Synergy Prediction,” (in eng), IEEE/ACM Trans Comput Biol Bioinform, vol. 19, no. 4, pp. 2334–2344, Jul-Aug 2022, doi: 10.1109/tcbb.2021.3086702.

[23] Q. Wang, X. Liu, and G. Yan, “Predicting effective drug combinations for cancer treatment using a graph-based approach,” Synthetic and Systems Biotechnology, vol. 10, no. 1, pp. 148–155, 2025/03/01/ 2025, doi: 10.1016/j.synbio.2024.09.003.

[24] C. Pilotto Heming et al., “P-Glycoprotein Drives Glioblastoma Survival and Chemotherapy Resistance: Potential as a Promising Liquid Biopsy Biomarker,” The American Journal of Pathology, vol. 195, no. 11, pp. 2131–2142, 2025/11/01/ 2025, doi: 10.1016/j.ajpath.2024.12.004.

[25] L. Wang and A. Sharma, “SERDs: a case study in targeted protein degradation,” (in eng), Chem Soc Rev, vol. 51, no. 19, pp. 8149–8159, Oct 3 2022, doi: 10.1039/d2cs00117a.

[26] Y. Huang, D. Jiang, M. Sui, X. Wang, and W. Fan, “Fulvestrant reverses doxorubicin resistance in multidrug-resistant breast cell lines independent of estrogen receptor expression,” (in eng), Oncol Rep, vol. 37, no. 2, pp. 705–712, Feb 2017, doi: 10.3892/or.2016.5315.

[27] P. Giannakakou et al., “A common pharmacophore for epothilone and taxanes: Molecular basis for drug resistance conferred by tubulin mutations in human cancer cells,” Proceedings of the National Academy of Sciences, vol. 97, no. 6, pp. 2904–2909, 2000/03/14 2000, doi: 10.1073/pnas.040546297.

[28] H. Ikeda et al., “Combination treatment with fulvestrant and various cytotoxic agents (doxorubicin, paclitaxel, docetaxel, vinorelbine, and 5-fluorouracil) has a synergistic effect in estrogen receptor-positive breast cancer,” (in eng), Cancer Sci, vol. 102, no. 11, pp. 2038–42, Nov 2011, doi: 10.1111/j.1349-7006.2011.02050.x.

[29] B. Biersack, B. Nitzsche, and M. Höpfner, “Histone deacetylases in the regulation of cell death and survival mechanisms in resistant BRAF-mutant cancers,” Cancer Drug Resistance, vol. 8, p. 6, 2025, doi: 10.20517/cdr.2024.125.

[30] D. W. Ball et al., “Trametinib with and without pazopanib has potent preclinical activity in thyroid cancer,” (in eng), Oncol Rep, vol. 34, no. 5, pp. 2319–24, Nov 2015, doi: 10.3892/or.2015.4225.

[31] P. A. Harris et al., “Discovery of 5-[[4-[(2,3-Dimethyl-2H-indazol-6-yl)methylamino]-2-pyrimidinyl]amino]-2-methyl-benzenesulfonamide (Pazopanib), a Novel and Potent Vascular Endothelial Growth Factor Receptor Inhibitor,” Journal of Medicinal Chemistry, vol. 51, no. 15, pp. 4632–4640, 2008/08/01 2008, doi: 10.1021/jm800566m.

[32] T. Żołek, A. Mazurek, and I.P. Gruzinski, “ n ilico tu ies of ovel Vemurafenib erivatives as Kinase Inhibitors,” Molecules, vol. 28, no. 13, p. 5273 doi: 10.3390/molecules28135273.

[33] R. Kurzrock et al., “A Phase I Trial of the VEGF Receptor Tyrosine Kinase Inhibitor Pazopanib in Combination with the MEK Inhibitor Trametinib in Advanced Solid Tumors and Differentiated Thyroid Cancers,” (in eng), Clin Cancer Res, vol. 25, no. 18, pp. 5475–5484, Sep 15 2019, doi: 10.1158/1078-0432.Ccr-18-1881.

[34] B. Biersack, L. Tahtamouni, and M. Höpfner, “Role and Function of Receptor Tyrosine Kinases in BRAF Mutant Cancers,” Receptors, vol. 3, no. 1, pp. 58–106 doi: 10.3390/receptors3010005.

[35] S. Taurin, M. Nimick, L. Larsen, and R. J. Rosengren, “A novel curcumin derivative increases the cytotoxicity of raloxifene in estrogen receptor-negative breast cancer cell lines,” (in eng), Int J Oncol, vol. 48, no. 1, pp. 385–98, Jan 2016, doi: 10.3892/ijo.2015.3252.

[36] J. Dinić et al., “Repurposing old drugs to fight multidrug resistant cancers,” Drug Resistance Updates, vol. 52, p. 100713, 2020/09/01/ 2020, doi: 10.1016/j.drup.2020.100713.

[37] J. Kowal, D. Ni, S. M. Jackson, I. Manolaridis, H. Stahlberg, and K. P. Locher, “Structural Basis of Drug Recognition by the Multidrug Transporter ABCG2,” Journal of Molecular Biology, vol. 433, no. 13, p. 166980, 2021/06/25/ 2021, doi: 10.1016/j.jmb.2021.166980.

[38] M. Minocha, V. Khurana, B. Qin, D. Pal, and A. K. Mitra, “Co-administration strategy to enhance brain accumulation of vandetanib by modulating P-glycoprotein (P-gp/Abcb1) and breast cancer resistance protein (Bcrp1/Abcg2) mediated efflux with m-TOR inhibitors,” (in eng), Int J Pharm, vol. 434, no. 1-2, pp. 306–14, Sep 15 2012, doi: 10.1016/j.ijpharm.2012.05.028.

[39] L. S. Zheng et al., “Vandetanib (Zactima, ZD6474) antagonizes ABCC1- and ABCG2-mediated multidrug resistance by inhibition of their transport function,” (in eng), PLoS One, vol. 4, no. 4, p. e5172, 2009, doi: 10.1371/journal.pone.0005172.

[40] K. Nishikawa, K. Kusama, H. Ekimoto, and K. Takahashi, “[Combination effects of etoposide with other antitumor drugs in vitro and in vivo],” (in jpn), Gan To Kagaku Ryoho, vol. 16, no. 12, pp. 3739–45, Dec 1989.

[41] V. Méresse et al., “Combined continuous infusion etoposide with high-dose cyclophosphamide for refractory neuroblastoma: a phase II study from the Société Française d’Oncologie Pédiatrique,” (in eng), J Clin Oncol, vol. 11, no. 4, pp. 630–7, Apr 1993, doi: 10.1200/jco.1993.11.4.630.

[42] J. Wang, X. Zhao, and L. Qiu, “Drug-induced self-assembled nanovesicles for chloroquine to sensitize MDR tumors to mitoxantrone hydrochloride,” Colloids and Surfaces B: Biointerfaces, vol. 245, p. 114358, 2025/01/01/ 2025, doi: 10.1016/j.colsurfb.2024.114358.

[43] A. L. Liang et al., “Chloroquine increases the anti-cancer activity of epirubicin in A549 lung cancer cells,” (in eng), Oncol Lett, vol. 20, no. 1, pp. 53–60, Jul 2020, doi: 10.3892/ol.2020.11567.

[44] S.V. Madhunapantula, A. Sharma, R. Gowda, and G.P. Robertson, “ entification of lyco en synthase kinase 3α as a therapeutic target in melanoma,” (in eng), Pigment Cell Melanoma Res, vol. 26, no. 6, pp. 886–99, Nov 2013, doi: 10.1111/pcmr.12156.

[45] F. Musumeci, A. Cianciusi, I.D. Aostino, G. Grossi, A. Carbone, and S. Schenone, “ ynthetic Heterocyclic erivatives as Kinase Inhibitors Tested for the Treatment of Neuroblastoma,” Molecules, vol. 26, no. 23, p. 7069 doi: 10.3390/molecules26237069.

[46] B. Wu, and Q. Wang. “DiffGraphTrans: a differential attention-based approach for extracting meaningful features of drug combinations.” Learning Meaningful Representations of Life (LMRL) Workshop at ICLR 2025. 2025. openreview.net/pdf?id=llfM9B2QML

[47] Q. Wang,, Z. Zhou, and G. Yan. “A Hypergraph-Based Model for Predicting Potential Drug Combinations in Cancer Therapy.” Interdisciplinary Sciences: Computational Life Sciences (2025): 1–11. doi: 10.1007/s12539-025-00779-3

